# Lewy pathology accumulates in swollen corticostriatal synapses in α-synucleinopathies

**DOI:** 10.1101/2025.08.27.672679

**Authors:** Alice Prigent, Michael X. Henderson

**Affiliations:** Department of Neurodegenerative Science, Van Andel Institute, Grand Rapids, MI 49503

**Keywords:** alpha-synuclein, putamen, corticostriatal, synapses, VGLUT1, Parkinson’s disease, dementia

## Abstract

Lewy body diseases (LBD), including Parkinson’s disease (PD), Parkinson’s disease dementia (PDD), and dementia with Lewy bodies (DLB), are neurodegenerative disorders characterized by the accumulation of misfolded α-synuclein in the form of Lewy pathology. A hallmark of these diseases is degeneration of the nigrostriatal pathway, resulting in loss of dopaminergic input to the striatum and consequent motor dysfunction. Lewy pathology is present in many regions outside the substantia nigra, and cortical Lewy pathology is the best correlate of cognitive decline in individuals that develop dementia. In addition, there is a high burden of neuritic Lewy pathology in the putamen, though the neuronal origin of this pathology is unclear. In the current study, we quantified the burden of Lewy pathology in the putamen across the spectrum of LBDs. Immunohistochemistry was used to quantify the Lewy burden in the putamen, cingulate and frontal cortices of 9 controls and 24 LBD cases. Even in PD cases without dementia, we observed a nearly complete striatal dopaminergic denervation among LBDs. Consistent with this denervation, most α-synuclein pathology did not co-localize with dopaminergic terminals, but was instead enriched in excitatory, VGLUT1-positive terminals. This enrichment in glutamatergic terminals was associated with swollen axons, but not with overall loss of VGLUT1 terminals. These findings suggest that Lewy pathology accumulates at excitatory corticostriatal synapses prior to overt synaptic degeneration and could contribute to cognitive decline in LBDs.

## INTRODUCTION

Parkinson’s disease (PD) is a progressive neurodegenerative disorder characterized primarily by motor symptoms such as bradykinesia, rigidity, tremor, postural instability, but also by a broad spectrum of non-motor symptoms including autonomic dysfunction, sleep disturbances, and cognitive impairment^1–4^. Central to PD and related disorders is the accumulation of misfolded α-synuclein protein aggregates, known as Lewy pathology, in vulnerable brain regions^5,6^. Over time, up to 80% of people with PD develop cognitive decline sufficient to warrant a diagnosis of Parkinson’s disease dementia (PDD)^7^. Dementia with Lewy bodies (DLB) shares overlapping clinical and pathological features with PDD^8,9^, but is distinguished primarily by the timing of symptom onset: in DLB, cognitive symptoms emerge before or within one year of motor symptoms, whereas in PDD, dementia develops after a longer duration of established motor disease^10–13^.

Lewy pathology was initially staged in 2003 by Braak and colleagues, who identified a characteristic spatiotemporal progression initially emerging in the olfactory structures and lower brainstem before ascending through midbrain regions such as the substantia nigra (SNpc) and eventually involving limbic and neocortical areas^14^. However, in 2009, Beach and colleagues proposed the Unified Staging System for Lewy Body Disorders to better categorize cases where Lewy pathology is limited to the olfactory bulb or follows a limbic-predominant pathway that initially does not involve the brainstem. This system classifies Lewy pathology stages as: 1) olfactory bulb only, 2) brainstem predominant or limbic predominant, 3) brainstem and limbic, and finally 4) neocortical^15^. This anatomical trajectory correlates broadly with the clinical trajectory of the disease: early non-motor symptoms, including hyposmia and autonomic dysfunction, correspond to pathology in peripheral and lower brainstem regions, while dopaminergic neuronal loss in the SNpc underlies the emergence of cardinal motor features^4,15–17^. In later stages, the involvement of neocortical regions is associated with cognitive decline and neuropsychiatric symptoms^18^, highlighting the multifaceted clinical impact of progressive α-synuclein aggregation throughout the nervous system.

Degeneration of the SNpc neurons leads to loss of dopaminergic input to the putamen, a key structure within the basal ganglia^19,20^. This loss of dopamine disrupts basal ganglia circuitry, resulting in impaired motor control^21–23^. The putamen plays a crucial role not only in motor learning and execution but also in cognitive functions^24^. It receives dense dopaminergic projections from the SNpc, as well as glutamatergic inputs from cortical regions and the thalamus, and GABAergic inputs from local collaterals as well as projections from GPe^25–27^. In addition to the loss of dopaminergic tone in putamen, this region also accumulates abundant Lewy neurites^28^. Interestingly, the burden of Lewy pathology in the putamen is even greater in PDD, DLB, and Alzheimer’s disease (AD), compared to PD alone, suggesting a relation of this pathology to dementia^29–31^. However, the precise origin of Lewy neurites in the putamen and their direct contribution to symptoms such as dementia remain unclear.

Presynaptic enrichment of Lewy pathology and synaptic dysfunction are hypothesized to be an early phenomenon in PD^32–34^. Recent studies have shown that α-synuclein aggregates can accumulate at dopaminergic synapses in the putamen prior to terminal loss, supporting a synaptic origin for pathology progression^35^. However, whether Lewy pathology similarly affects other striatal afferents, particularly glutamatergic and GABAergic inputs, and how this relates to cortical Lewy burden, remains less explored in Lewy body disorders (LBD).

In the current study, we sought to determine which specific presynaptic inputs to the putamen—corticostriatal glutamatergic (VGLUT1-positive), or GABAergic (VGAT-positive)—are preferentially affected by Lewy pathology across PD, PDD, and DLB. Additionally, we investigated whether α-synuclein accumulation at these terminals is associated with structural changes, such as synaptic swelling, which may contribute to synaptic dysfunction. We examined the burden of cortical and striatal Lewy pathology in 8 controls, 9 idiopathic PD, 10 idiopathic PDD and 9 idiopathic DLB. Co-immunofluorescence was used to assess Lewy pathology colocalization with dopaminergic (TH), cortical glutamatergic (VGLUT1) and GABAergic (VGAT) terminals in the putamen across all groups. All disease groups showed a nearly complete denervation of dopaminergic terminals, so only a minimal proportion Lewy pathology was located in these terminals. Inhibitory VGAT-positive terminals were also only rarely associated with Lewy pathology. In contrast, the majority of Lewy pathology was associated with excitatory VGLUT1-positive terminals. The majority of VGLUT1 inputs to the putamen come from corticostriatal projections. Indeed, cortical Lewy pathology load was positively associated with putamen Lewy pathology. Interestingly, across LBDs, swollen VGLUT1 puncta were observed, and these swollen puncta were colocalized with Lewy aggregates. Despite the enrichment Lewy pathologyVGLUT1-positive terminals, there was no overall reduction in the number of terminals. Taken together, our results suggest that Lewy pathology accumulation at cortical excitatory synapses may precede detectable synaptic degeneration in the putamen and could contribute to the cortical propagation of Lewy pathology and progression to cognitive decline.

## RESULTS

### Case demographics

A total of 36 cases were included in this study: 8 controls (CTRL), 9 Parkinson’s disease (PD), 10 Parkinson’s disease with dementia (PDD) and 9 dementia with Lewy bodies (DLB). The sex, postmortem interval (PMI), age of death and disease duration are shown in (**Table S1**). Despite efforts to match for age, control cases were significantly younger than the PD donors (61 vs 77 years, p=0.001) and the PDD donors (61 vs 72 years, p=0.018). No significant differences were observed compared to the DLB donors (61 vs 70 years, p=0.055) (**Fig. 1A**). Disease duration was significantly shorter in DLB cases compared to PD and PDD cases (**Fig. 1B**). The postmortem interval, representing the time elapsed between represents the time between death and freezing, was similar across groups (**Fig. 1C**). For most of the groups, the male /female ratio was higher (CTRL: F= 37.5%; PDD: F= 0%; DLB: F= 11%) except for the PD (F= 57%, **Fig. 1D**).

**Figure 1.**
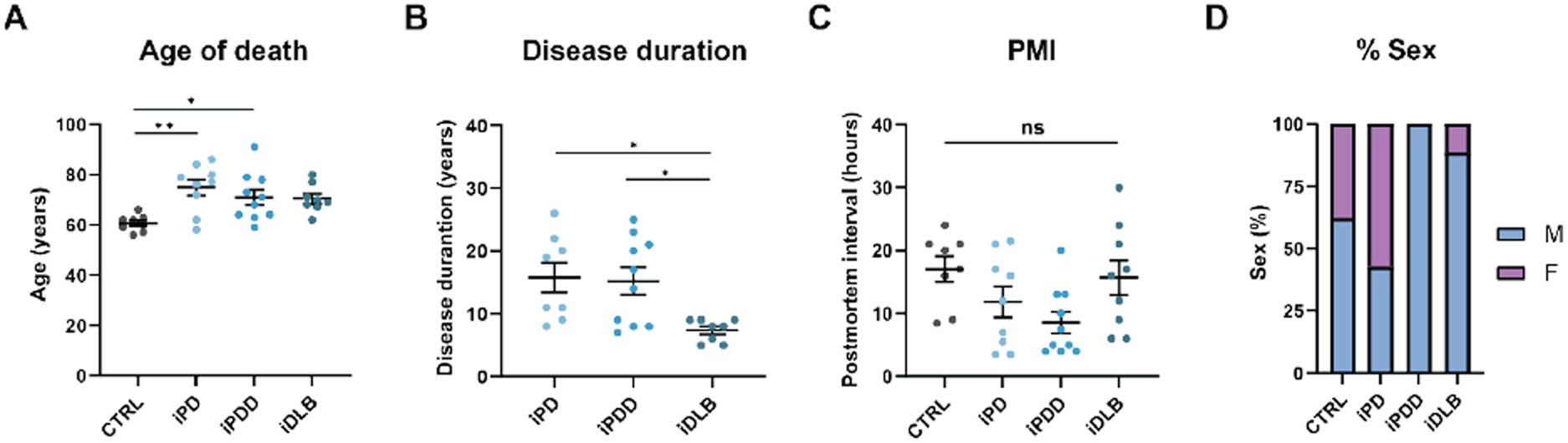
Demographic and clinical characteristics of study cohort. **A**. Age at death of all cases. **B**. Disease duration. **C**. Post-mortem interval (PMI). **D**. Percentages of total group represented by each sex. Male (M) are shown in blue and female (F) in violet. Bars represent mean ± S.E.M. with individual values plotted. One-way ANOVA; Tukey’s multiple comparison test. *p < 0.05, **p < 0.01, ns= non significant.

### Striatal Lewy pathology burden is associated with cortical Lewy pathology across α-synucleinopathies

The burden of α-synuclein pathology in the putamen is not well-characterized and is not included in general staging systems^14^. However, in a previous study focused on the relationship between lipid metabolites and protein pathologies, we observed abundant Lewy neurites in the putamen of PD, PDD, and DLB cases^31^. In the current study, we aimed to further understand which projection neurons bear this pathology. Further, we had observed some larger inclusions in the putamen and aimed to understand if these were somatic Lewy bodies. The chosen subset of cases displayed elevated pathology in the putamen, cingulate, and frontal cortex in PD, PDD, and DLB compared to controls (**Fig. 2A, 2B**). As reported in other studies^14,36^, pathology in the cingulate cortex is more severe than in frontal cortex.

We next investigated whether the burden of Lewy pathology in the putamen was related to cortical pathology, since one of the major inputs to the striatum is cortico-striatal projections. Lewy pathology levels in the frontal were strongly correlated with those in the putamen across all groups (**Fig. 2C**).

**Figure 2.**
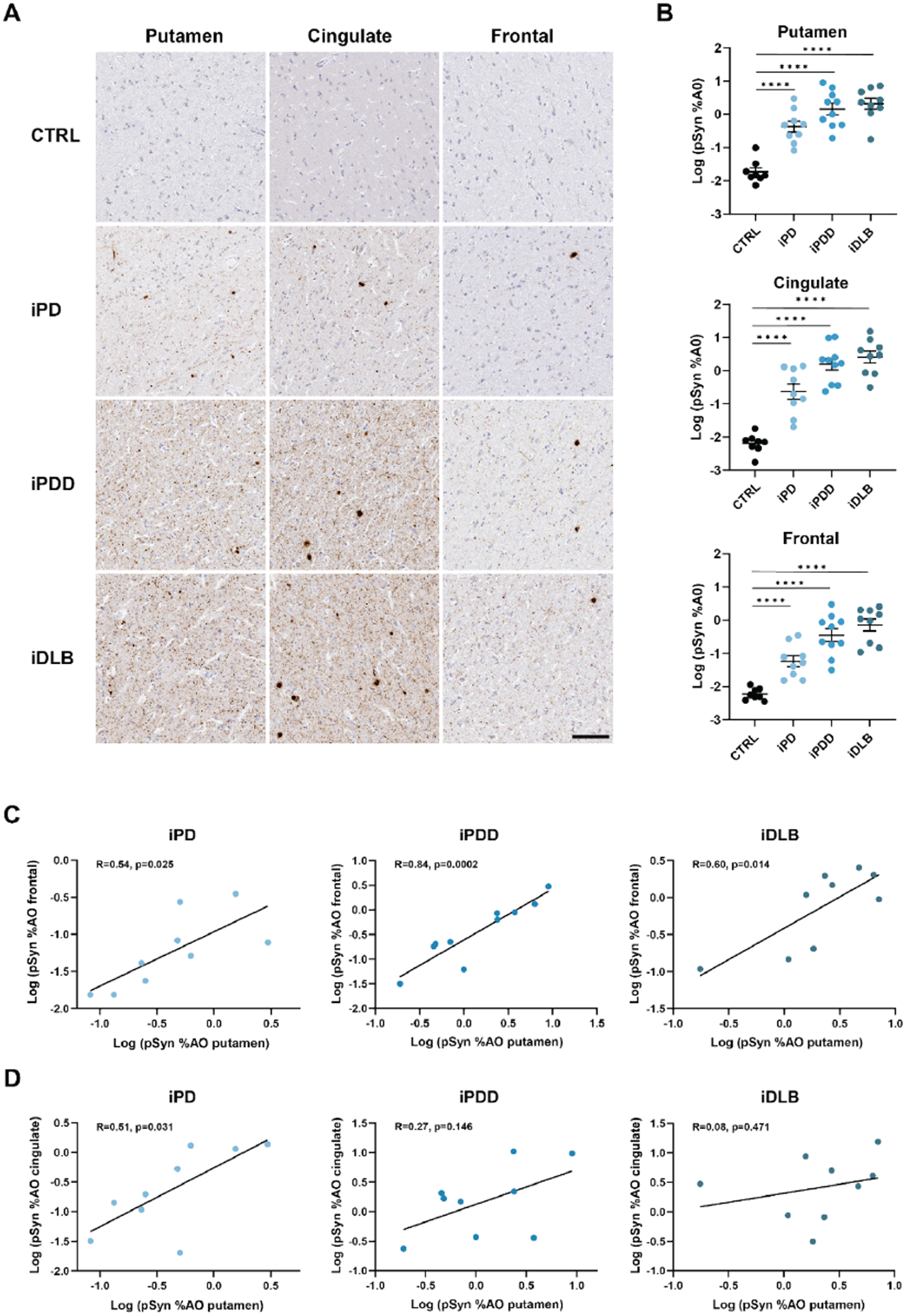
Quantitative Lewy pathology and neuropathological correlations in α-synucleinopathies. **A.** Representative staining of pS129 α-synuclein (pSyn) in the putamen, cingulate, and frontal cortex of iPD, iPDD and iDLB cases. Scale bar = 100μm. **B**. Quantification of pSyn levels in putamen, cingulate and frontal cortex. Bars represent mean ± S.E.M. with individual values plotted. One-way ANOVA; Tukey’s multiple comparison test. *p < 0.05, **p < 0.01, ***p < 0.001, ****p < 0.0001. **C**. Correlation between levels of pSyn in putamen and frontal cortex. **D**. Correlation between pSyn levels in putamen and cingulate cortex. Lines represent linear regression line of best-fit.

In contrast, pSyn levels in the cingulate were only positively correlated with the putamen in the PD group, while no significant correlation was observed for the PDD or DLB groups (**Fig. 2D**). This lack of significance may be due to the overall high pathology burden in dementia cases, as across all cases, putamen pathology was correlated with cingulate pathology (**Fig. S1**). Together, these findings suggest that the neuritic pathology in the putamen could represent cortico-striatal projections.

### Tyrosine hydroxylase projections are degenerated across LBDs and co-localize minimally with pSyn pathology

Dopaminergic neurons from the substantia nigra project to the putamen and are severely impacted in LBDs. Based on staining, these vulnerable dopaminergic neurons appear to undergo axonal dieback, and eventually neurons are lost^14,37,38^. Therefore, we first analyzed the expression of dopaminergic terminals in the putamen using immunofluorescence labelling for tyrosine hydroxylase (TH) (**Fig. 3A**). The putamen showed severe denervation in LBD cases (**Fig. 3A, 3B**), similar to previous reports^19,28^. No significant differences were observed between α-synucleinopathies, suggesting that all patients were at an advanced stage of the the disease, resulting in nearly complete striatal dopaminergic denervation at the time of death.

**Figure 3.**
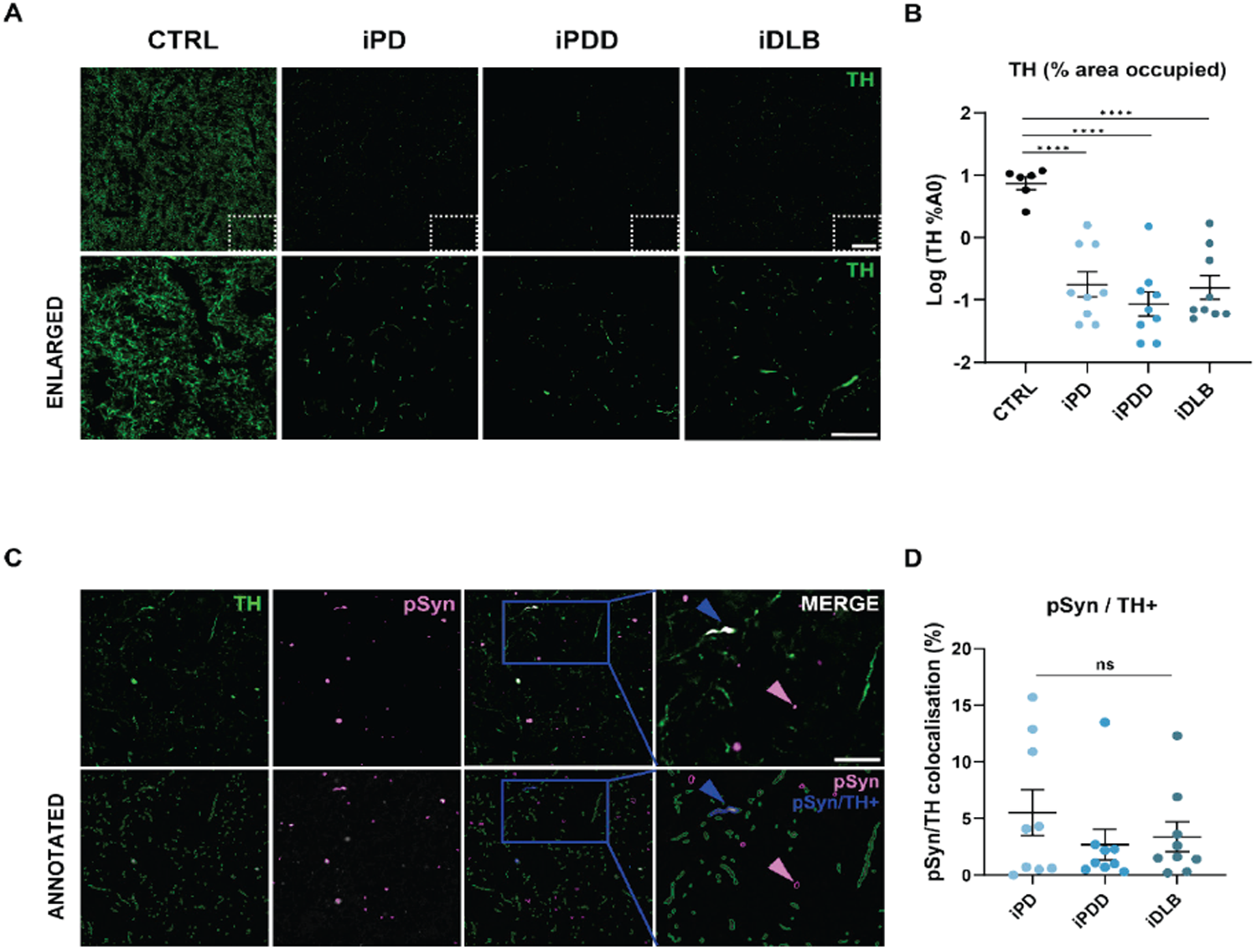
Tyrosine hydroxylase projections are degenerated across LBDs and co-localize minimally with pSyn pathology. **A.** Representative images of tyrosine hydroxylase (TH) immunofluorescence in the putamen of CTRL, iPD, iPDD and iDLB cases. Scale bar = 50 μm (first row), Scale bar (Enlarged images) = 20 μm (second row). **B**. Quantification of percentage of TH area occupied across all groups. Graph bars represent mean ± S.E.M. with individual values plotted. One-way ANOVA; Tukey’s multiple comparison test. *p < 0.05, **p < 0.01, ***p < 0.001, ****p < 0.0001. **C**. Representative images of pSyn colocalization with TH fibers in the putamen (first row). Annotated images of object-based colocalization (second row): pSyn/TH+ (blue arrow) and pSyn not colocalized (magenta arrow). Scale bar = 20 μm. **D**. Quantification of the percentage of pSyn objects colocalizing with TH across all groups. Graph bars represent mean ± S.E.M. with individual values plotted. One-way ANOVA; Tukey’s multiple comparison test. ns= non-significant.

To determine whether striatal Lewy pathology was in the remaining TH fibers, we used an object-based colocalization approach to determine the portion of pSyn objects within TH-positive fibers. Confocal images co-stained for pSyn and TH were analyzed using CellProfiler software, where pSyn aggregates colocalizing with TH-positive fibers were identified as pSyn/TH^+^objects (**Fig. 3C**). Only a small fraction (2-6%) of pSyn aggregates colocalized with TH-positive fibers, with no significant differences observed across groups (**Fig. 3D**). Overall, these results suggest that, in our cohort group, by the late stages of α-synucleinopathies, dopaminergic terminals in the putamen are already largely degenerated, and pSyn pathology accumulates predominantly outside the remaining dopaminergic fibers.

### Striatal Lewy pathology is enriched in VGLUT1 presynaptic terminals

The putamen receives dense excitatory input from the cerebral cortex, particularly from motor and associative regions. Disruption of this corticostriatal connectivity, driven by α-synuclein pathology and synaptic dysfunction, may contribute to the cognitive decline and dementia^18^. The putamen is predominantly composed of GABAergic neurons and these neurons send collateral inputs within the putamen^26,27^. To assess potential alterations in excitatory and inhibitory synaptic inputs, we quantified glutamatergic cortical and GABAergic terminals using immunofluorescence labeling for VGLUT1 and VGAT, respectively, followed by object-based analysis (**Fig. 4A**). No significant differences in the VGLUT1 and VGAT levels were observed among groups (**Fig. 4B, 4C**). The number of pSyn objects were elevated in disease groups compared to controls, and significantly increased in DLB compared to controls and PD. (**Fig. 4D**).

**Figure 4.**
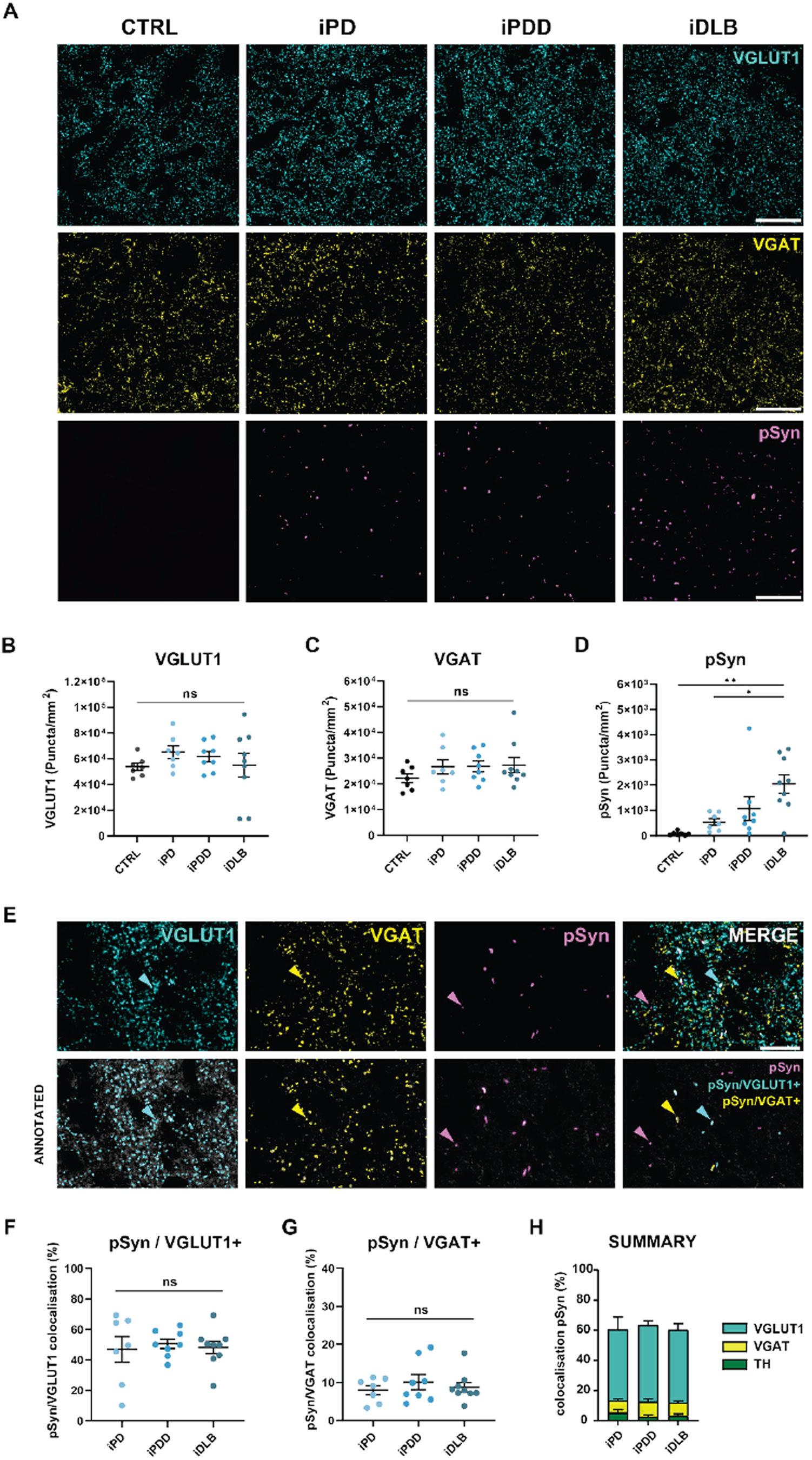
Striatal Lewy pathology is enriched in VGLUT1 presynaptic terminals. **(A)** Representative staining of VGLUT1, VGAT and pSyn in the putamen of iPD, iPDD and iDLB cases. Scale bar =50μm. **(B–D)** Quantification of VGLUT1 **(B)**, VGAT **(C)**, and pSyn **(D)** puncta per mm^2^ across groups. **(E)** Representative images of pSyn colocalization with presynaptic markers VGLUT1 and VGAT (top row) and annotated object-based colocalization (bottom row): pSyn/VGLUT1^+^(cyan arrows), pSyn/VGAT^+^(yellow arrows), and pSyn without colocalization (magenta arrows). Scale bar = 20 μm. **(F–G)** Quantification of the percentage of pSyn objects colocalized with VGLUT1 **(F)** and VGAT **(G)** across groups. **(H)** Summary of pSyn colocalization expressed as cumulative percentage in stacked bar format. Graph bars represent mean ± SEM, with individual data points shown. Statistical analysis performed using one-way ANOVA with Tukey’s multiple comparisons test. *p < 0.05; **p < 0.01; ***p < 0.001; ****p < 0.0001; ns = not significant.

Using the same object-based colocalization approach applied in the TH analysis, we next quantified the proportion of pSyn aggregates colocalized with VGLUT1-positive or VGAT-positive synaptic terminals, defined as pSyn^+^/VGLUT1^+^and pSyn^+^/VGAT^+^, respectively (**Fig. 4E**). We observed that pSyn aggregates mostly (∼50%) colocalized with VGLUT1-positive terminals **(Fig. 4F)**. In contrast, colocalization with VGAT-positive terminals was minimal (∼9%) (**Fig. 4G**). Collectively, these findings indicate that approximately half of the Lewy pathology within the putamen is localized to glutamatergic presynaptic terminals, yet this enrichment is not accompanied by a reduction ofVGLUT1-positive terminals.

### VGLUT1 presynaptic terminals containing pSyn pathology are swollen

Many of the α-synuclein inclusions in the putamen were large enough to be small Lewy bodies. However, upon co-staining, these larger inclusions co-localized with larger VGLUT1-positive puncta **(Fig. 5A**). Recent studies have shown that Lewy pathology enrichment at presynaptic terminals may contribute to their swelling, dysfunction and eventual loss^35,39^. To assess potential changes in synaptic morphology, we analyzed the size of VGLUT1-positive presynaptic terminals with (VGLUT1/pSyn+) and without (VGLUT1) pSyn pathology using CellProfiler software (**Fig. 5A**). In the segmented images (second row), we observed enlarged VGLUT1 synapses colocalizing with pSyn, highlighted by orange arrows (**Fig. 5A**). We observed a 2-fold increase in the size of VGLUT1 presynaptic terminals with pSyn compared to VGLUT1 without, but no significant difference observed between LBD groups (**Fig. 5B**). These results demonstrate that while α-synuclein inclusions in the putamen are associated with dopaminergic terminal loss, glutamatergic terminals remain, but become dystrophic, potentially preceding eventual synapse and neuron loss.

**Figure 5.**
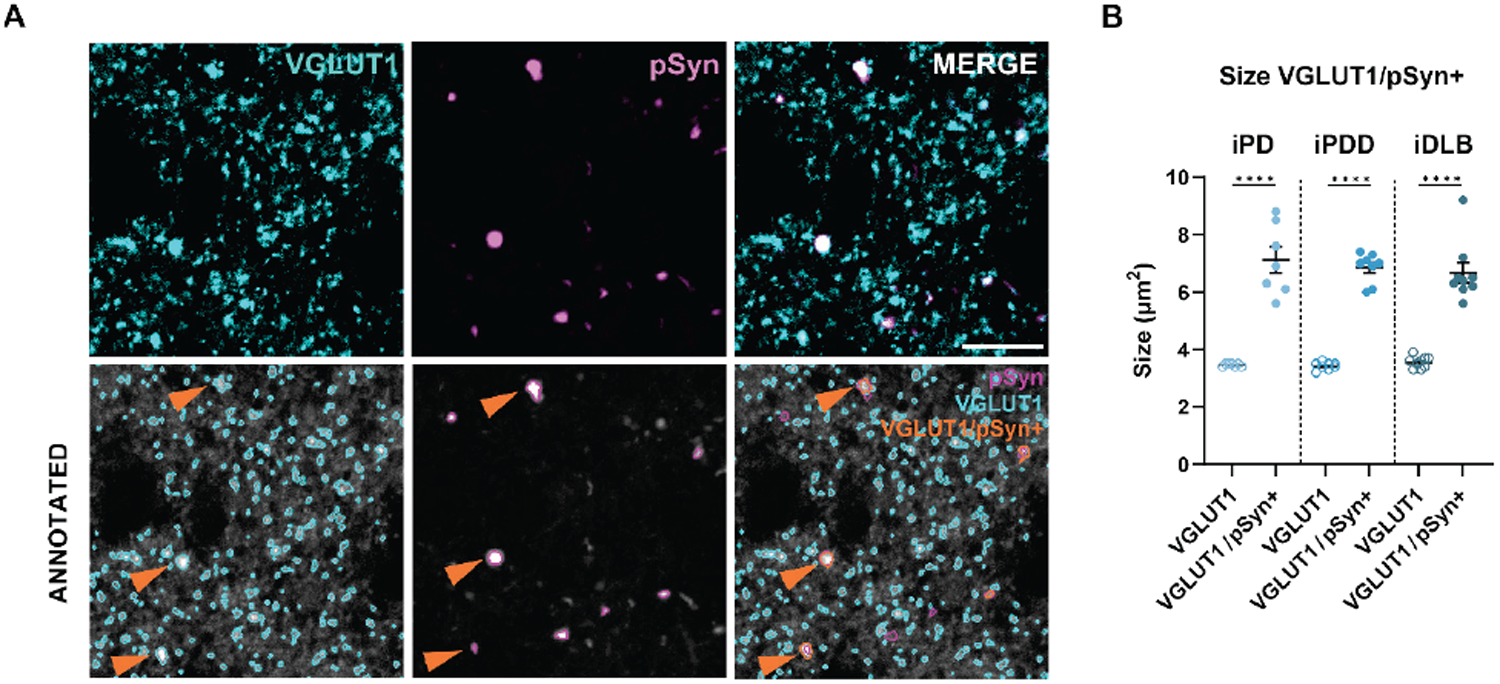
Enlarged VGLUT1 terminals contain Lewy pathology. **A.** Representative images of pSyn colocalization with enlarged presynaptic marker VGLUT1 in the putamen (first row). Annotated images: VGLUT1 maker (cyan) and pSyn (magenta), enlarged pSyn/VGLUT1+ (orange arrow), Scale bar =20μm. **B**. Quantification of the size of presynaptic markers VGLUT1 with (VGLUT1/pSyn+) or without pSyn (VGLUT1) colocalized. Graph bars represent mean ± S.E.M. with individual values plotted. Unpaired Student’s t-test for each group: ****p < 0.0001.

## DISCUSSION

In this study, we aimed to understand which neurons projecting to the putamen bear neuritic Lewy pathology across LBDs and how that pathology relates to structural changes in synapses. We found that 1) dopaminergic denervation was advanced across all groups, minimizing the extent of Lewy pathology in those terminals, 2) Lewy pathology was selectively enriched in putamen glutamatergic VGLUT1-positive synaptic terminals, 3) Lewy pathology was associated with swelling of terminals but not overall presynaptic terminal loss in the putamen, and 4) Lewy pathology in the putamen correlated with Lewy pathology in cortical regions. Taken together, our findings highlight a circuit-specific vulnerability of corticostriatal synapses and suggest that presynaptic Lewy pathology may contribute to cortical disease progression and cognitive impairment in LBDs.

α-Synuclein is a presynaptic protein involved in synaptic vesicle trafficking and neurotransmitter release, that plays a central role in the pathogenesis of Lewy body disorders^32,40,41^. Misfolded α-synuclein can propagate trans-synaptically in model systems, spreading from cell-to-cell along anatomically and functionally connected networks^42,43^. This prion-like property enables α-synuclein to propagate through vulnerable brain circuits, driving progressive synaptic dysfunction, neuronal death, and disconnection within and between specific cell populations^42,44^. However, despite evidence implicating α-synuclein in synaptic pathology, the enrichment of pSyn in specific cell-type terminals remains largely unexplored. Our data confirm and extend prior observations^14^ of a rostro-caudal gradient of Lewy pathology burden, showing significant pSyn accumulation in the putamen across PD, PDD, and DLB, with progressively higher levels in the cingulate and frontal cortices paralleling the emergence and severity of cognitive symptoms.

The degeneration of nigrostriatal dopaminergic projections is a defining pathological feature of PD and related α-synucleinopathies. Clinically, it is estimated that by the time of PD diagnosis, 50–70% of striatal dopaminergic terminals have already been lost^17,37,45^. This early and severe denervation is closely linked to the emergence of motor symptoms. In line with these observations, we observe a pronounced loss of tyrosine hydroxylase (TH)-immunoreactive fibers within the putamen across all disease groups. The absence of significant differences between PD, PDD, and DLB suggests that all cases were at an advanced disease stage, including those clinically diagnosed as PD. As expected, we find that only a small proportion of pSyn aggregates colocalized with residual TH-positive fibers in the putamen, and this proportion is comparably low across all diagnostic groups. These findings indicate that by late disease stages, Lewy neurites in the putamen are largely in non-dopaminergic neurons. Our results are consistent with recent study that shows pSyn enrichment at dopamine transporter (DAT)-positive terminals in incidental Lewy body disease, but not in PD^35^. This suggests Lewy pathology shifts from dopaminergic to glutamatergic terminals in the putamen as disease progresses.

α-Synuclein is enriched in excitatory synapses^46,47^ and has Lewy pathology is localized to layer 5 intratelencephalic (IT) neurons in postmortem PD cortex^48^, highlighting glutamatergic as one of the neuron types vulnerable to developing Lewy pathology. However, the extent to which pathological α-synuclein accumulates in corticostriatal synapses across LBD is only recently being explored. A key finding of this study is that approximately half of all pSyn aggregates in the putamen colocalize with VGLUT1-positive glutamatergic terminals across PD, PDD, and DLB, while colocalization with VGAT-positive GABAergic terminals was consistently minimal. This selective synaptic enrichment supports a model wherein excitatory corticostriatal projections are a major site of α-synuclein aggregation, potentially reflecting local uptake of misfolded α-synuclein by cortical axon terminals within the striatum. Such a mechanism would be consistent with trans-synaptic propagation models, in which α-synuclein released by degenerating dopaminergic terminals or other affected neurons in the striatum is internalized by corticostriatal axons and retrogradely transported to cortical region, contributing to the spread of pathology and cortical dysfunction in LBD. Interestingly, the proportion of VGLUT1-positive pSyn aggregates did not differ significantly across disease groups (∼50%), which may reflect a plateau effect in late-stage disease. The origin of the remaining pSyn signal that does not colocalize with either VGLUT1-or VGAT-positive terminals remains to be determined. One plausible interpretation, is that these aggregates reside within excitatory axons that lack a presynaptic specialization within the imaging volume. Alternatively, a subset of pSyn aggregates may originate from neuronal populations not assessed in this study, such as thalamostriatal or brainstem-derived inputs. However, considering the established presynaptic enrichment of α-synuclein and its preferential association with excitatory terminals, it is likely that a significant proportion of these non-colocalized aggregates represent glutamatergic terminals lacking detectable synaptic markers.

We observed that VGLUT1-positive presynaptic terminals colocalizing with pSyn aggregates were significantly enlarged, which could reflect a functional as well as a morphological disruption. Indeed, neuritic α-synuclein aggregates can disrupt synaptic vesicle trafficking, leading to their clustering and swelling synapses and eventually their pruning or loss^34,49^. This is in line with previous studies reporting that swollen synapses and synaptic dysfunction are an early event in PD, DLB and PD models in response to pSyn aggregates^35,39,50^. Furthermore, dopaminergic terminal loss in the putamen occurs before the degeneration of neuronal cell bodies in the SNpc^19^, emphasizing the early role of synaptic pathology in disease progression.

Corticostriatal glutamatergic transmission, plays a critical role in the manifestation of motor and cognitive symptoms observed in LBD. A study using immunofluorescence labeling and synaptic density quantification, showed no changes in VGLUT1 expression in incidental PD or across PD disease stage (Braak 1 to 6) in the striatum^35^. Additionally, a semi-quantitative analysis by western blot and immunoautoradiography showed an increase in VGLUT1 in the putamen associated with decrease VGLUT1 expression level in frontal and temporal cortices^51^. In agreement with the two human studies, we observe a non-significant trend toward increased VGLUT1- and VGAT-positive puncta in the putamen in PD and PDD compared to controls, suggesting that the overall abundance of cortical and inhibitory inputs to the putamen remains preserved at the terminal level despite clinically associated dementia symptoms. Nevertheless, variation in the level of VGLUT1 expression level within the DLB group demonstrates that some cases may have substantial excitatory terminal loss. PDD and DLB brains harbor more amyloid-β plaques in the striatum than PD and striatal plaques are associated with dementia severity^52,53^. These observations raise the possibility that in DLB, compared to PD and PDD, striatal VGLUT1 loss could be driven in part by amyloid-β pathology.

This study has several limitations. First, our analysis focused primarily on VGLUT1-positive glutamatergic terminals, which selectively label cortical input to the striatum. As a result, we did not assess thalamostriatal projections, which are predominantly VGLUT2-positive^54,55^. This was due to poor VGLUT2 antibody performance in human tissue. Therefore, we cannot exclude the possibility that a substantial fraction of pSyn aggregates may also localize to VGLUT2 terminals, particularly given the fact that consequent Lewy pathology is found in the thalamus of PD patients^56^. Second, while we VGLUT1-positive terminals were not changed in PD and PDD, this apparent resilience may reflect compensatory sprouting or an increase in structurally intact but functionally impaired synapses. Because our approach did not include assessment of postsynaptic specializations or synaptic function, we cannot determine whether these terminals are functionally competent or merely structurally retained. Finally, our cases represent later stages of disease, though they are in agreement with other work that has assessed a broader range of Braak stages^35^.

Together, these findings support a model of selective vulnerability within corticostriatal circuits, where early synaptic α-synuclein aggregation may contribute to cortical disease progression and cognitive impairment in LBD.

## AVAILABILITY OF DATA AND MATERIALS

The data that support the findings of this are available upon request.

## ACKNOWLEDGEMENTS

We would like to thank the patients and families who participated in this research, without whom this study would not have been possible. We would like to thank Terry Schuck and Brian Alfaro with assistance in tissue collection and the Van Andel Institute

## AUTHORS’ CONTRIBUTIONS

A.P. designed experiments, performed experiments, analyzed results and wrote the paper. M.X.H designed the experiments and wrote the paper. All authors have reviewed and approved the paper,

### List of abbreviations

AD: Alzheimer’s disease
CTRL: Control
DAT: Dopamine transporter
DLB: Dementia with Lewy bodies
LB: Lewy body
LBD: Lewy body diseases
PD: Parkinson’s disease
PDD: Parkinson’s disease dementia
PMI: Postmortem interval
pS129: Phospho-serine 129
pSyn: Phospho-α-synuclein
SNpc: Substantia nigra, pars compacta
TH: Tyrosine hydroxylase
VGAT: Vesicular GABA transporter
VGLUT1: Vesicular glutamate transporter 1
VGLUT2: Vesicular glutamate transporter 2

## FUNDING

This work was funded in part by the Van Andel Institute.

Pathology and Biorepository Core for tissue sectioning (RRID:SCR_022912).

## MATERIAL AND METHODS

### Human brain tissue

A total of 36 brains were used in this study: 8 controls, 9 PD, 10 PDD and 9 DLB. All brain tissues were obtained from the Center of Neurodegenerative Disease Research (CNDR) at the University of Pennsylvania. The study protocol was approved by the University of Pennsylvania ethics committee and written informed consent was obtained from next-of-kin. Frozen tissue was briefly thawed and transferred to 10% neutral buffered formalin (NBF) for overnight fixation. The fixed tissue was embedded in paraffin after 24 h for further histological examination. Primary pathology to estimate % area occupied of pS129 α-synuclein was conducted on these tissues and previously published^31^.

### Immunofluorescence

Slides were de-paraffinized with 2 sequential 5 min washes in xylenes, followed by 1 min washes in a descending series of ethanols: 100%, 100%, 95%, 80%, 70%. Slides were then incubated in deionized water for one min prior and transferred to the BioGenex EZ-Retriever System where they were incubated in antigen unmasking solution (Vector Laboratories; Cat# H3300) and microwaved for 15 min at 95 °C. Slides were allowed to cool for 20 min at room temperature, washed in running tap water for 10 min and then blocked for 1 h at RT in 0.1 M Tris/2% fetal bovine serum (FBS). Primaries antibodies were incubated in 0.1 M Tris/2% FBS in a humidified chamber overnight at 4°C. The following antibodies were used: mouse monoclonal anti-pS129 α-synuclein (81A) (1:1000; Biolegend, Cat# MMS-5091, RRID: AB_2564891), guinea pig anti-VGLUT1 (1:1000; Synaptic Systems, Cat#135 138, RRID: AB_887875), rabbit polyclonal anti-VGAT (1:1000; synaptic system, Cat#131 002, RRID: AB_887871), mouse monoclonal anti-tyrosine hydroxylase (1:1000; Sigma-Aldrich, Cat#T2928, RRID: AB_477569). Primary antibodies were rinsed off with 0.1 M tris for 5 min and then incubated with the appropriate secondary antibody for 1 hour at room temperature: Goat anti-Rabbit IgG (H+L), Alexa Fluor™ 488 (1:1000; Thermo Fisher Scientific, Cat # A-11008; RRID:AB_143165), Goat anti-Guinea Pig IgG (H+L), Alexa Fluor™ 568 (1:1000; Thermo Fisher Scientific, Cat # A-11075; RRID:AB_2534119), Goat anti-

Mouse IgG1, Alexa Fluor™ 568 (1:1000; Thermo Fisher Scientific, Cat # A-21124; RRID:AB_2535766), Goat anti-Mouse IgG2a, Alexa Fluor™ 647 (1:1000; Thermo Fisher Scientific, Cat # A-21241; RRID:AB_2535810). Secondaries antibodies were rinsed off with 0.1M tris for 5min and then treated with Sudan black B (Thermo Fisher Scientific, Cat #190160250) for 45 seconds. Slides were rinsed off and then mounted with coverglass in ProLong gold with DAPI (Invitrogen, Cat#P36931).

Fluorescence images were acquired using a Nikon A1plus-RSi laser-scanning confocal microscope (Nikon Instruments) using a 60x oil immersion objective. For the synaptic markers, four images of 500 μm x 500 μm were captured from the putamen of each patient. For the tyrosine hydroxylase analysis, three large images of 1 mm each were imaged.

### Colocalization object-based and object size quantification

Fluorescence images were imported into CellProfiler software (CellProfiler v4.2.6). After applying the RescaleIntensity module, primary objects were detected using the speckles (synaptic makers) and neurites (pSyn) features. Synaptic objects with diameters smaller than 0.4 μm and pSyn objects smaller than 1 μm were excluded from analysis.Child and parent objects were assigned and the parent with the most overlap was considered colocalized with child object. pSyn objects colocalizing with VGLUT1-positive terminals and VGAT-positive terminals presynaptic markers were defined as pSyn/VGLUT1+ and pSyn/VGAT+, respectively. The number of total, colocalized objects and object size (µm^2^) were then determined.

For TH analysis, primary objects were detected using the neurite feature. pSyn objects colocalizing with TH are defined as pSyn/TH+. Area occupied by TH, pSyn colocalized objects were then determined.

### Statistical analysis

GraphPad Prism software version 9.3.1 (GraphPad Software Inc., La Jolla, CA, USA) was used for pair-wise statistical analysis. The data shown in this study are mean ± standard error of the mean (SEM). Unpaired Student t-test was used to determine differences between two groups. For comparison of multiple groups, a Brown-Forsythe test was first applied to test if variances were significantly different. If group variances were not different, a one-way ANOVA was applied with Tukey’s multiple comparison test to determine differences between any groups. Linear regressions and correlation coefficients were all calculated in GraphPad Prism.

