## Supplemental Figures for "Lewy pathology accumulates in swollen corticostriatal synapses in α-synucleinopathies"

| Group | Sex | PMI | Age of Death | Disease duration |
|---|---|---|---|---|
| **Control** | Female | 21 | 63 | N/A |
| **Control** | Male | 8.5 | 62 | N/A |
| **Control** | Male | 24 | 57 | N/A |
| **Control** | Female | 21 | 56 | N/A |
| **Control** | Male | 16 | 62 | N/A |
| **Control** | Male | 20 | 66 | N/A |
| **Control** | Female | 9 | 60 | N/A |
| **Control** | Male | 17 | 59 | N/A |
| **iPD** | Female | 21 | 80 | 11 |
| **iPD** | Male | 16 | 86 | 26 |
| **iPD** | Female | 3.5 | 77 | 11 |
| **iPD** | Male | 3.5 | 62 | 9 |
| **iPD** | Male | 7 | 84 | 19 |
| **iPD** | Female | 17 | 79 | 20 |
| **iPD** | Female | 5.5 | 72 | 22 |
| **iPD** | Female | 12 | 76 | N/A |
| **iPD** | Male | 21.5 | 58 | 8 |
| **iPDD** | Male | 7.5 | 78 | 25 |
| **iPDD** | Male | 4 | 79 | 9 |
| **iPDD** | Male | 4 | 64 | 17 |
| **iPDD** | Male | 5 | 73 | 14 |
| **iPDD** | Male | 20 | 63 | 23 |
| **iPDD** | Male | 5 | 71 | 7 |
| **iPDD** | Male | 10 | 59 | 20 |
| **iPDD** | Male | 13 | 91 | 8 |
| **iPDD** | Male | 4 | 68 | 8 |
| **iPDD** | Female | 13 | 64 | 21 |
| **iDLB** | Male | 24 | 71 | 9 |
| **iDLB** | Male | 30 | 77 | 8 |
| **iDLB** | Male | 16 | 80 | 9 |
| **iDLB** | Male | 9 | 71 | 9 |
| **iDLB** | Male | 12 | 68 | 8 |
| **iDLB** | Male | 17 | 62 | N/A |
| **iDLB** | Male | 6 | 70 | 6 |
| **iDLB** | Female | 6 | 69 | 5 |
| **iDLB** | Male | 21 | 67 | 5 |

**Table S1. Patients' demographics.**
PMI: postmortem interval

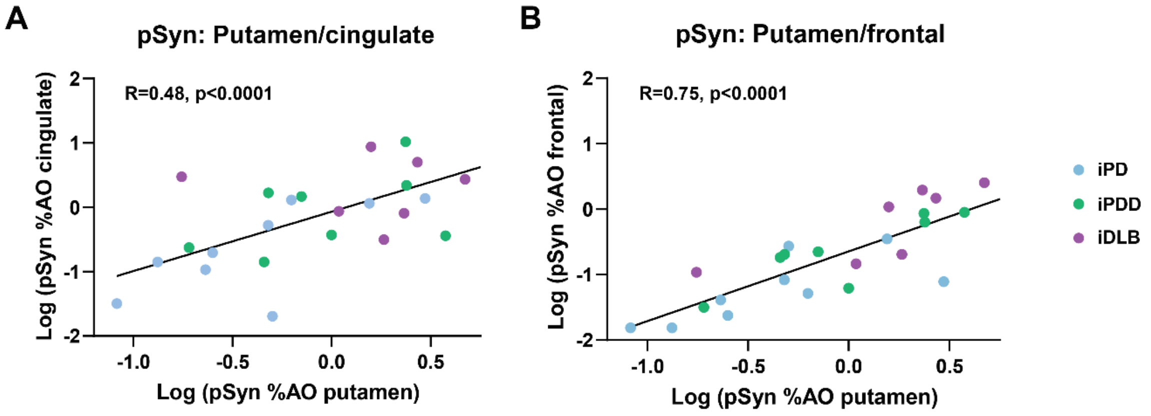


**Figure S1: Neuropathological correlations in α-synucleinopathies**. **A.** Correlation between pSyn levels in the putamen and cingulate cortex. **B.** Correlation between pSyn levels in the putamen and frontal cortex. Lines represent linear regression line of best-fit.
